# Role of DNA methylation dynamics in desiccation and salinity stress responses in rice cultivars

**DOI:** 10.1101/558064

**Authors:** Mohan Singh Rajkumar, Rama Shankar, Rohini Garg, Mukesh Jain

**Author notes:** MSR **RS:** **RG:** **MJ:**.

## Abstract

DNA methylation is an epigenetic mark that controls gene expression in response to internal and environmental cues. In this study, we sought to understand the role of DNA methylation in response to desiccation and salinity stresses in three rice cultivars (IR64, stress-sensitive; Nagina 22, drought-tolerant and Pokkali, salinity-tolerant) via bisulphite sequencing. We identified DNA methylation patterns in different genomic/genic regions and analysed their correlation with gene expression. Methylation in CG context within gene body and methylation in CHH context in distal promoter regions were positively correlated with gene expression. However, methylation in other sequence contexts and genic regions was negatively correlated with gene expression. DNA methylation was found to be most dynamic in CHH context under stress condition(s) in the rice cultivars. The expression profiles of genes involved in *de-novo* methylation were correlated with methylation dynamics. Hypomethylation in Nagina 22 and hypermethylation in Pokkali in response to desiccation and salinity stress, respectively, were correlated with higher expression of abiotic stress response related genes. Our results suggest an important role of DNA methylation in abiotic stress responses in rice in cultivar-specific manner. This study provides useful resource of DNA methylomes that can be integrated with other data to understand abiotic stress response in rice.

**Highlight:** Bisulphite sequencing revealed single base resolution DNA methylation, and cultivar-specific differential methylation patterns and correlation with gene expression that control desiccation and salinity stress response in the rice cultivars.

## Introduction

Abiotic stress is one of the most unpredictable contraventions resulting in crop loss and imposes major concerns in food security. Global warming and adverse environmental conditions result in lower crop yield. Rice is one of the most important cereal crops serving as primary source of dietary carbohydrates in more than half of world population. Drought and salinity stresses affect growth and yield of rice (Bartels *et al*., 2005; Singh *et al*., 2018; Peethambaran *et al*., 2018). Unlike animals, plants are sessile in nature and responses to abiotic stress are associated with activation of gene regulatory networks and pathways involved in stress response (Shinozaki *et al*., 2007; Zeller *et al*., 2009). The role of transcription factors and signal transduction components in adaptive stress responses has been revealed (Xiong *et al*., 2001; Mizoi *et al*., Zhu *et al*., 2016; Khan *et al*., 2018). Epigenetic modifications associated with chromatin re-organization can also play significant role in stress responses (Bruce *et al*., 2007; Probst *et al*., 2015; Garg *et al*., 2015; Neto *et al*., 2017).

DNA methylation is one of most commonly found epigenetic marks. In plants, methylated cytosines are found in three different sequence contexts [CG, CHG and CHH (where H = A, C or T)]. Maintenance of DNA methylation after each round of DNA replication in daughter stands is mediated by METHYLTRANSERASE 1 (MET1) in CG context and maintenance in CHG context is mediated via CHROMOMETHYLASE 3 (Lindroth *et al*., 2001; Kankel *et al*., 2003; Law *et al*., 2010). De-novo methylation in CHH context is governed by small RNA dependent and independent pathways via DOMAINS REARRANGED METHYLTRANSFERASE 1/2 (DRM1/2) and CHROMOMETHYLASE 2 (CMT2), respectively (Cao *et al*., 2002; Slothkin *et al*., 2009; Mosher *et al*., 2010; Stroud *et al*., 2014).

The possible role of DNA methylation in abiotic stress response has been reported in model and crop plants (Chinnusamy *et al*., 2009; Lang *et al*., 2010). Drought induced methylation differences have been identified in sensitive and tolerant cultivars via methylation-sensitive restriction digestion and whole genome bisulphite sequencing in different plants (Wang *et al*., 2011; Liang *et al*., 2014; Garg *et al*., 2015). Interestingly, methylation dynamics under drought stress has been found extensively in CHH context in both susceptible and tolerant cultivars in *Populus trichocarpa* (Liang *et al*., 2014). However, methylation differences between drought-tolerant and susceptible cultivars under control conditions were mostly found in CG context in rice (Garg *et al*., 2015). Likewise, methylation dynamics under salinity stress have also been investigated in susceptible and tolerant cultivars in different plants using various approaches, and methylation changes under the stress conditions were found to be genotype-specific (Wang *et al*., 2014; Al-Lawati *et al*., 2016; Lu *et al*., 2017). Interestingly, methylation changes induced in response to abiotic stress condition(s) reversed to its original methylation status after stress recovery (Wang *et al*., 2011). However, the plants exposed to those abiotic stress(s) were found to adapt more successfully to adverse environmental conditions in subsequent exposures and epigenetic memory was transmitted to subsequent generations (Boyko *et al*., 2010; Bilichak *et al*., 2016).

Earlier, we reported DNA methylation patterns and differences between rice cultivars, including stress-sensitive IR64, drought-tolerant Nagina 22 and salinity-tolerant Pokkali (Garg *et al*., 2015). To further understand the role of DNA methylation in desiccation and salinity stress responses, we sequenced DNA methylomes of rice seedlings exposed to these stresses and compared with that of control conditions. We analyzed the extent of DNA methylation in different sequence contexts at whole genome level. Methylation density and patterns of DNA methylation in protein coding genes and transposable elements (TEs) were revealed. The correlation between DNA methylation density and gene expression levels was analysed under stress conditions in the rice cultivars. We identified differentially methylated regions (DMRs) under stress conditions and their correlation with differential gene expression was revealed. The possible role of DNA methylation in regulation of sets of genes involved in abiotic stress response was also analyzed. Overall, we demonstrated the role of DNA methylation in abiotic stress responses in cultivar-specific manner in rice.

## Materials and Methods

### Plant materials and genomic DNA isolation

We analyzed three rice cultivars with contrasting response to abiotic stresses, including tolerant to desiccation (Nagina 22), tolerant to salinity (Pokkali) and sensitive to both stress conditions (IR64). Two-weeks old hydroponically grown rice seedlings were treated with desiccation stress in IR64 and Nagina 22 by keeping them between folds of tissue papers for 3 h. Salinity stress was given by keeping seedlings in 200 mM NaCl solution for 3 h. The control plants were kept in water for the same duration as described in the previous study (Garg *et al*., 2015). Stress treated and control seedlings were harvested, snap frozen in liquid nitrogen and stored at −80 °C till further use. Genomic DNA was isolated using Qiagen DNeasy Minikit (Qiagen) as per manufacturer’s instructions. Genomic DNA was quantified using Qubit Fluorimeter (Life Technologies) and purity of DNA was verified by estimating absorbance ratio at 260/280 and 260/230 wavelengths using Spectrophotometer (Nanodrop) and integrity of the DNA was verified by resolving into agarose gel.

### Whole genome bisulphite sequencing

To prepare library for bisulphite sequencing, we fragmented the genomic DNA of all the tissue samples to an average size of 100-300 bp via sonication (Covaris, Massachusetts, USA). The genomic fragments were end repaired and TrueSeq-methylated adaptors were ligated to their ends. Adaptor ligated genomic fragments were treated with sodium bisulphite as described in previous study (Garg *et al*., 2015). Library preparation and sequencing were performed to generate 90 nt long reads in paired-end mode with sufficient sequencing depth (>30x) via HiSeq-2000 platform (Illumina, San Diego, USA).

### Read alignment and identification of mCs

The adaptor sequences and low-quality reads were removed from the raw reads using NGSQC Toolkit (v2.3) at default parameters (Patel and Jain, 2012). The clonal reads were filtered out by mapping on the rice genome (MSU v7.0) using Bismark (v0.8) under default parameters in each sample (Krueger and Andrews, 2011). Efficiency of bisulphite conversion was estimated by mapping the high-quality filtered reads on rice chloroplast genome. More than >99% of the cytosine(s) in chloroplast genome were converted to thymine(s) indicating very high efficiency of bisulphite conversion in our experiments. The mCs in rice genome were identified based on the significance of ≤0.001 *p*-value and sequencing depth of ≥5 reads, as described in previous study (Garg *et al*., 2015). Methylation level was determined by estimating the percentage of reads giving methylation call at a particular cytosine site to all the reads in the sequencing data covering that site (Garg *et al*., 2015). Patterns of DNA methylation in rice genome were visualized via circos plot using a window size of 100 kb. Density of DNA methylation in genes/TEs and their 2 kb flanking regions was calculated using customized perl scripts.

### Identification of DMRs

We identified differential methylation under stress conditions as compared to the control condition for each cultivar within each 100 bp bin in the rice genome. The bins covered by ≥5 reads and containing at least 3 cytosine residues were considered for differential methylation analysis. The two bins with same genomic co-ordinates under control and stress conditions were analysed for detection of methylation level difference. Differentially methylated bins showing at least 20% methylation level difference with <0.01 *q*-value calculated using Fisher’s exact test followed by correction with Sliding Linear Model (SLIM) were determined as described in the previous studies (Garg *et al*., 2015, Bhatia *et al*., 2018). Consecutive differentially methylated bins (within a distance of 50 bp) were merged to identify DMRs and their distribution in different sequence contexts within gene body and 2 kb flanking regions was analyzed.

### Gene ontology (GO) enrichment analysis

The enrichment of gene ontology (GO) terms in the sets of DMR associated genes in different cultivars was analysed using BiNGO tool in Cytoscape (v3.7). The significantly enriched GO terms with *q*-value of at least ≤0.05 were identified for each given set of genes.

### Integration of DNA methylation and gene expression data

The correlation between DNA methylation and gene expression was determined by plotting methylation density of genes expressed at different levels in each sample. Based on FPKM values, genes expressed at very low (<1 FPKM), low (≥1-5 FPKM), moderate (≥5-25 FPKM) and high (>25 FPKM) levels were categorized. Methylation density in gene body, gene ends and flanking regions was estimated for these sets of genes expressed at varying levels. The correlation between differential methylation and differential gene expression (≥2 fold-change with <0.05 *q*-value) under desiccation/salinity stress as compared to control condition within same cultivar was analysed by estimating methylation level differences in different sequence contexts within gene body and flanking regions.

### Data availability

Bisulphite sequencing and RNA sequencing data reported/used in this study are available under GSE60288 and GSE60287 series accession numbers, respectively, at the Gene Expression Omnibus (GEO) database.

## Results and discussion

### Methylome profiling under desiccation and/or salinity stress

To understand the role of DNA methylation in desiccation and salinity stress responses, we profiled DNA methylomes in seedlings of IR64 (sensitive), drought-tolerant (Nagina 22) and salinity-tolerant (Pokkali) rice cultivars after the stress treatment(s) and compared with that of respective control condition. We analysed DNA methylation in IR64 under desiccation and salinity stresses, Nagina 22 under desiccation stress and Pokkali under salinity stress. Bisulphite sequencing allowed interrogating DNA methylation status of rice genome at single base resolution. We generated about (76-86 million) high-quality 90 bp long paired-end reads with >30x sequencing depth for each sample. A total of about 46-55 million uniquely mapped reads covered ∼88% of the rice genome and (81.5-82.8 %) of the total cytosine residues in each sample (Table S1).

We estimated percentage of methylcytosines (mCs) with respect to total cytosines in each sequence context that were covered in sequencing for each sample analysed. Interestingly, about half (46.7-49.52%) of the total cytosines in CG context were found to be methylated in each sample followed by CHG (28.36-30.72%) and CHH contexts (19.76-24.3%) (Fig. 1A). Methylation level of mCs in CG context (85.45-90.37%) was much higher than CHG (66.41-70.15%) and CHH (39.76-45.69%) contexts (Fig. 1B; Fig. S1). Previous studies have also shown that methylation levels are generally higher in CG context (Lister *et al*., 2008; Cokus *et al*., 2008; Garg *et al*., 2015; Hossain *et al*., 2017). It may be due to methylation maintenance in CG and CHG contexts that ensures methylation in the newly formed DNA strands after replication. Another reason for high methylation level in these sequence contexts may be due to KRYPTONITE mediated methylation in CHG context in deep heterochromatic regions associated with H3K9me2 followed by spreading of methylation in CG context (Du *et al*., 2014; Trejo *et al*., 2017). In contrast, de-novo methylation in CHH context is dependent on internal and/or environmental cues, which may be the possible reason for low methylation level in CHH context. Similar methylation levels were found in forward and reverse strands in all the samples analysed (Fig. S2) as reported in previous studies (Jones *et al*., 2007; Probs *et al*., 2009; Garg *et al*., 2015).

**Fig. 1.**
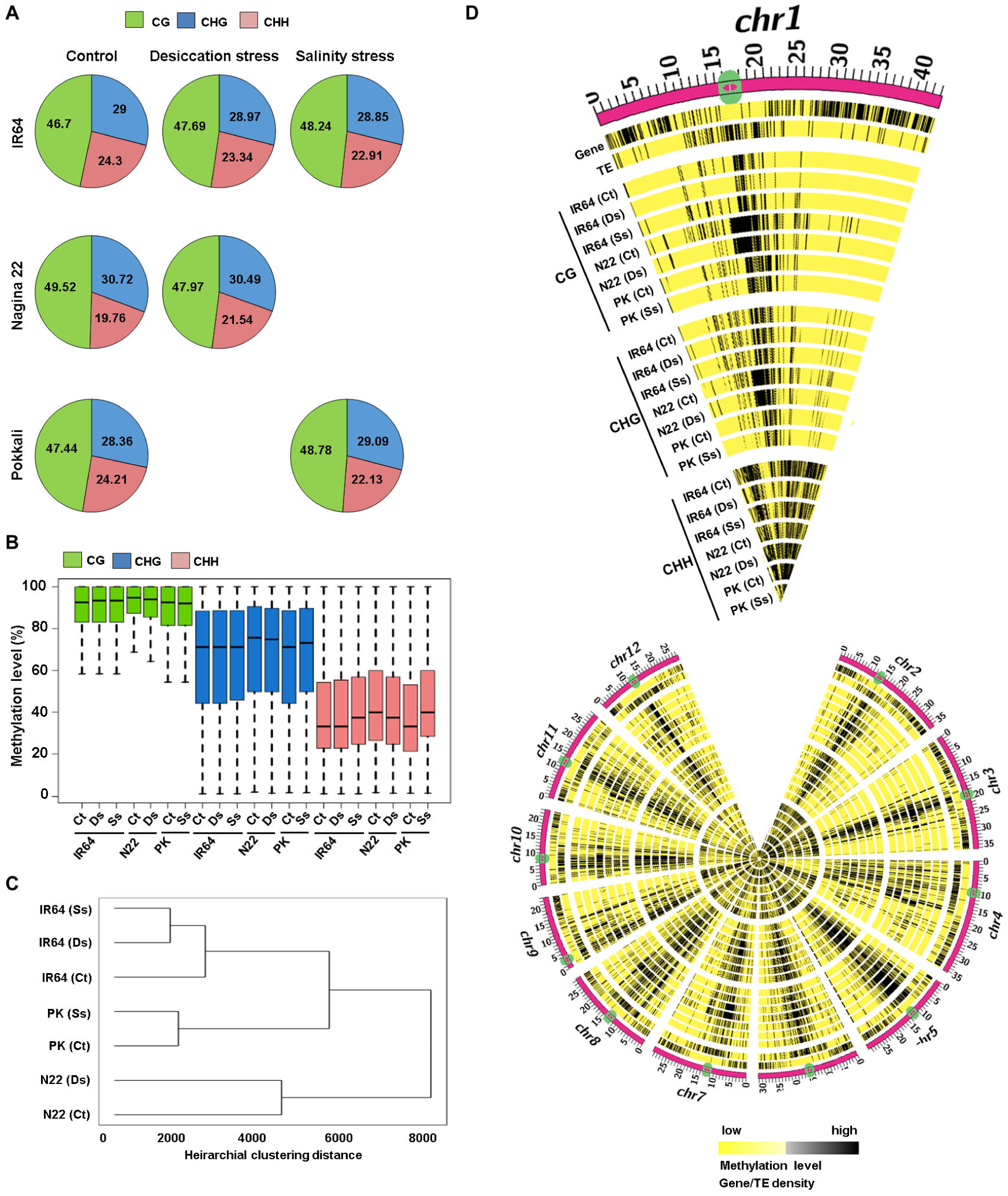
Methylation patterns in the rice cultivars under desiccation and salinity stresses. (A) Pie chart showing percentage of methylated cytosines in different sequence contexts under control and stress conditions in the three rice cultivars. (B) Methylation levels in different sequence contexts under control and stress conditions are shown in boxplot. (C) Dendrogram showing similarity/divergence of methylation patterns among the rice cultivars under control and stress conditions are shown. (D) Methylation density (number of methylated cytosines per 100 kb) in rice genome under control and stress conditions is shown via circos plot. Gene and TE density has also been shown in the two outermost circles. Ct, control; Ds, desiccation stress; Ss, salinity stress; N22, Nagina 22; PK, Pokkali.

To examine differences in patterns of DNA methylation in response to abiotic stress, we compared methylation levels between stress treated and control samples in each cultivar. Interestingly, methylation levels showed most variations in CHH context followed by CHG context in response to desiccation and salinity stresses in Nagina 22 and Pokkali, respectively (Fig. 1B). Decreased methylation levels in Nagina 22 under desiccation stress and increased methylation level in Pokkali under salinity stress were observed in both the sequence contexts. However, no obvious methylation level difference was observed in these sequence contexts under desiccation and salinity stress conditions in IR64 with the exception of increased methylation level detected in CHH context under salinity stress. In CG context, methylation levels were marginally higher under both stress conditions in IR64. In contrast, methylation levels in CG context were marginally lower under desiccation stress in Nagina 22 and under salinity stress in Pokkali. These results showed methylation level differences majorly in CHH context under stress conditions, suggesting its important role in determining abiotic stress response.

To understand correlation among different samples analysed, we performed clustering among control and stress treated samples of all the rice cultivars based on detected mCs. The control and stress treated sample(s) of the same cultivar clustered together, suggesting that methylome divergence between the cultivars is much higher than the methylome dynamics in response to abiotic stress within a cultivar (Fig. 1C). Further, to examine global methylation patterns, we analyzed the distribution of DNA methylation on the rice chromosomes in all the samples. Interestingly, pericentromeric and centromeric regions harbouring high density of transposable elements (TEs) were found to be extensively methylated in CG and CHG contexts in control and stress treated samples in the rice cultivars. In contrast, higher fraction of mCs in CHH context was detected in gene rich regions under control and stress conditions in all the cultivars, further suggesting the important role of CHH context DNA methylation in abiotic stress response (Fig. 1D).

### DNA methylation in protein coding genes and TEs

We estimated DNA methylation density within the body of protein coding genes and TEs, and their flanking regions in different sequence contexts in all the samples analysed. In general, methylation density in TEs was much higher than protein coding genes in all the sequence contexts. The methylation density at gene ends representing transcription start site (TSS) and transcription termination site (TTS) was much lower than their body and flanking regions in all the sequence contexts. Interestingly, decreased methylation density at TE ends was not observed in CG and CHG contexts, suggesting important role of DNA methylation in TE repression (Fig. 2). Another interesting difference in methylation patterns between genes and TEs was observed in CHH context. In genes, methylation density at proximal promoter regions (−500 bp) is significantly high in CHH context, but no such distinct methylation pattern was observed in TEs (Fig. 2).

**Fig. 2.**
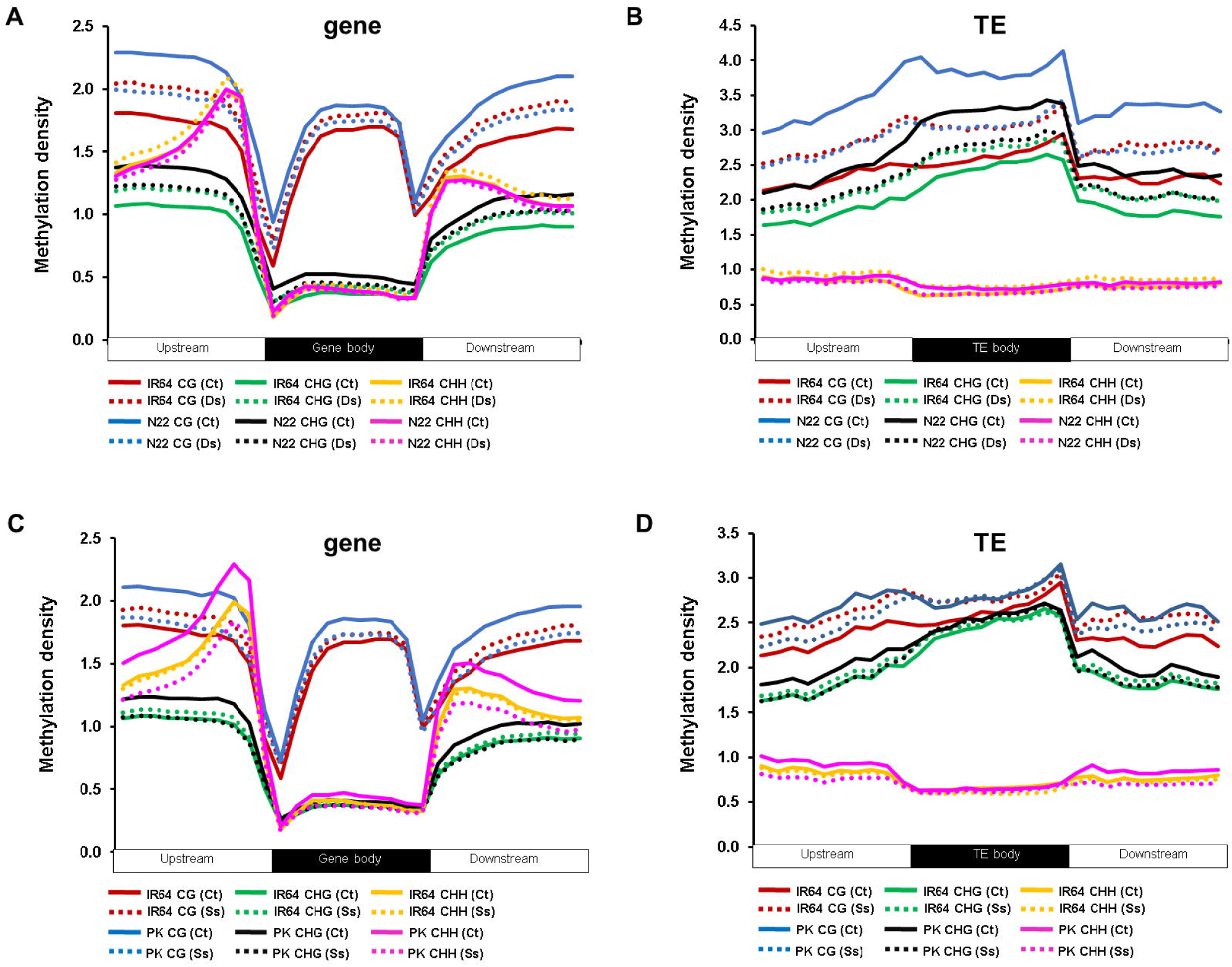
Methylation patterns in genes and transposable elements (TEs) under desiccation and/or salinity stresses in the rice cultivars. (A, B) Methylation density in IR64 and Nagina 22 is shown in genes (A) and TEs (B) in different sequence contexts under control and desiccation stress conditions. (C, D) Methylation density in genes (C) and TEs (D) in IR64 and Pokkali under control and salinity stress conditions is shown. Methylation density indicates number of methylcytosines (mCs) per 100 bp.

Next, we analysed differences in methylation density under stress condition(s) as compared to control in the rice cultivars. In Nagina 22, methylation density under desiccation stress was lower as compared to control condition in all the sequence contexts. In contrast, methylation density was higher under desiccation stress in IR64. The difference in methylation density was most evident in the flanking (−500 bp) regions (Fig. 2A). Interestingly, similar differential pattern of DNA methylation profiles between control and desiccation stress treated samples was detected in TEs in both sensitive and tolerant rice cultivars (Fig. 2B; Fig. S3A). Likewise, methylation density difference between control and salinity stress conditions in IR64 and Pokkali cultivars was analysed. Higher methylation density in IR64 and lower methylation density in Pokkali were observed in all the sequence contexts under salinity stress in both protein coding genes and TEs (Fig. 2C, D; Fig. S3B).

### Influence of DNA methylation on gene expression

To examine influence of DNA methylation on expression of protein coding genes, we analysed methylation of genes expressed at varying levels under control and stress conditions. The rice genes were classified into sets of genes based on their expression levels, including silent/very-low (<1 FPKM), low (≥1 to 5 FPKM), moderate (≥5 to 25 FPKM) and high (>25 FPKM) using transcriptome data from our previous study (Shankar *et al*., 2016). A positive correlation between methylation density in CG context of gene body and expression level was observed as reported in previous studies (Zilberman *et al*., 2007; Bewick *et al*., 2017). Interestingly, we observed a positive correlation between DNA methylation in CHH context in flanking regions (−500 bp) and gene expression. A positive correlation between DNA methylation in CHH context in distal promoter regions and TE expression in rice has been reported in previous studies (Zemach *et al*., 2010a; Zemach *et al*., 2010b). The mechanism of CHH methylation in promoter regions and enhanced gene expression is largely unknown. In contrast, an antagonistic correlation between DNA methylation in all other genic regions in CG and CHG contexts and gene expression was observed. However, most significant antagonistic correlation of gene expression was found with methylation at transcription start site (TSS) and transcription termination site (TTS) in all the sequence contexts, suggesting that methylation at TSS/TTS may repress gene expression (Fig. 3). These results suggest an important role of DNA methylation in determining expression levels of genes irrespective of different cutivar(s) and/or condition(s).

**Fig. 3.**
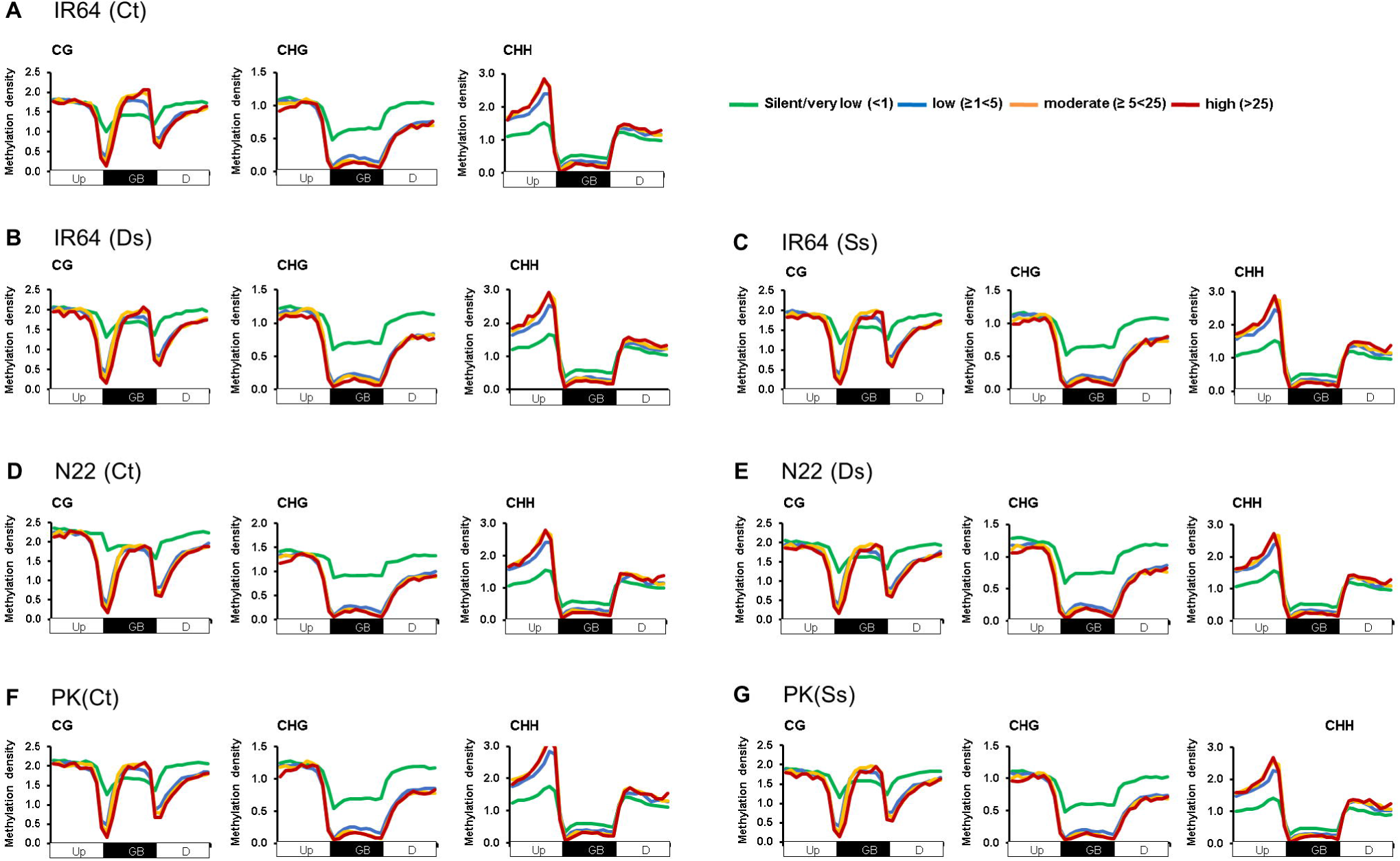
Influence of DNA methylation on gene expression. (A-G) Methylation density of genes expressed at different levels [silent/very-low (<1 FPKM), low (≥1 to 5 FPKM), moderate (≥5 to 25 FPKM) and high (>25 FPKM)] under control [IR64 (A), Nagina 22 (D), Pokkali (F)], desiccation stress [IR64 (B), Nagina 22 (E)] and salinity stress [IR64 (C), Pokkali (G)] conditions are shown. Up, upstream region, GB, gene body; D, downstream region.

### Differentially methylated regions under abiotic stress in rice cultivars

To study methylation dynamics under desiccation stress in IR64 and Nagina 22 rice cultivars, differentially methylated regions (DMRs) between stress and control conditions were identified for both the cultivars. In total, 2346 and 3013 DMRs representing 2162 and 2744 genes were detected in IR64 and N22, respectively, under desiccation stress. Interestingly, highest number of DMRs was found in CHH context (76.73-77.92%) followed by CG (15.33-17.22%) and CHG (4.86-7.93%) contexts in both the cultivars, suggesting an important role of CHH context DNA methylation in response to desiccation stress (Fig. 4A). Further, we analyzed hyper/hypomethylation under desiccation stress in different sequence contexts. Interestingly, about 80%, 78% and 71% of the total DMRs were found to be hypomethylated in CG, CHG and CHH contexts, respectively, under desiccation stress in Nagina 22. Number of hypermethylated DMRs in CG (59.41%) and CHH (53.83%) contexts were marginally higher, while number of hypomethylated DMRs in CHG context (55.26%) were lower under desiccation stress in IR64 (Fig. 4A). Next, we estimated methylation level differences under desiccation stress in both the cultivars. In IR64, fraction of hypermethylated DMRs in CG and CHG contexts was more than hypomethylated DMRs under desiccation stress. In contrast, much higher fraction of DMRs was found to be hypomethylated in all sequence contexts in Nagina 22 (Fig. 4B, C). These results suggest that hypomethylation may be associated with desiccation stress response in Nagina 22.

**Fig. 4.**
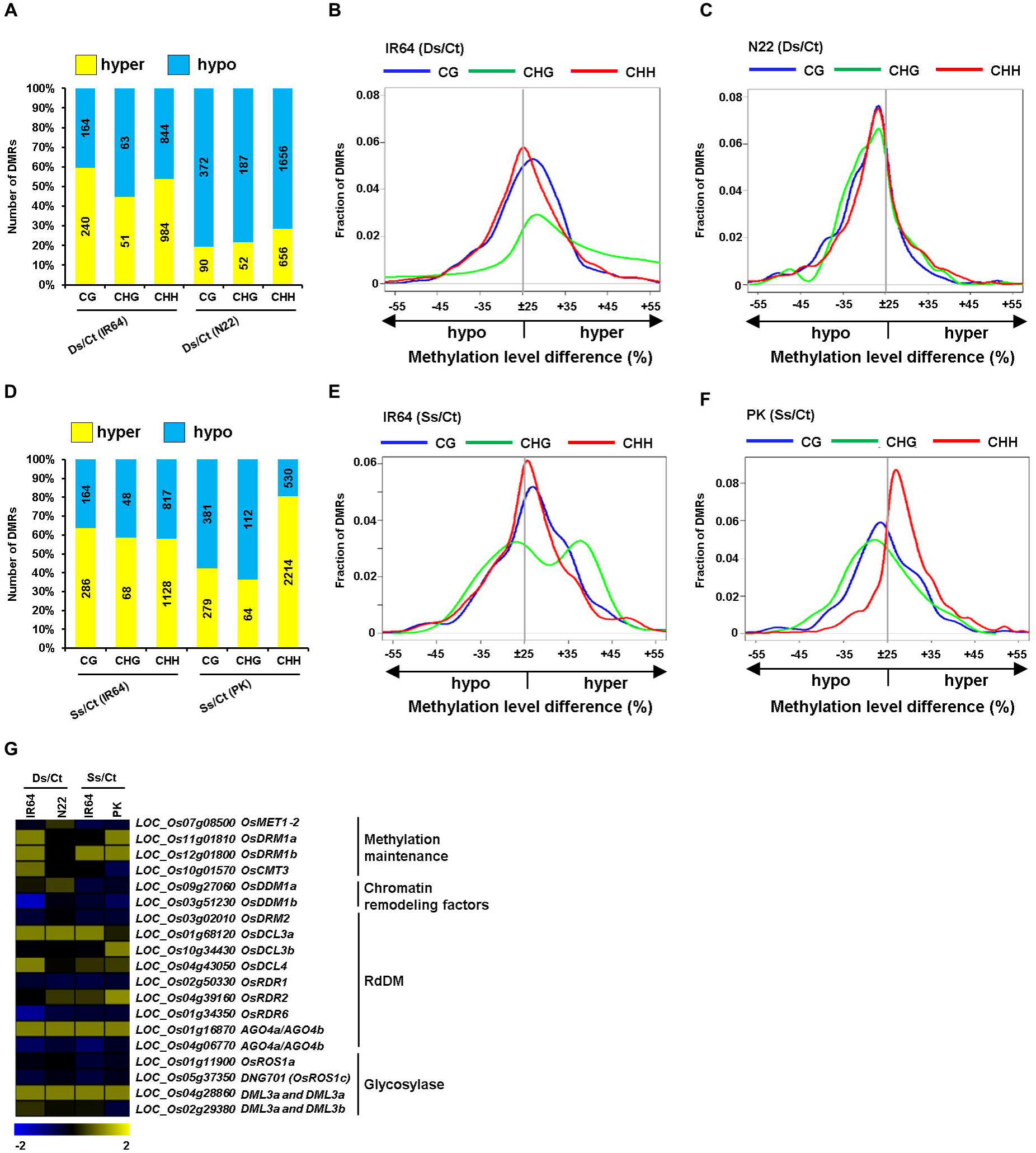
Methylation dynamics in rice cultivars under desiccation and/or salinity stresses. (A) Number of differentially methylated regions (DMRs) in different sequence contexts under desiccation stress in IR64 and Nagina 22 are shown. (B, C) Methylation level differences in different sequence contexts under desiccation stress in IR64 (B) and Nagina 22 (C) are shown via kernel density plots. (D-F) Number of DMRs in different sequence contexts under salinity stress in IR64 and Pokkali are shown (D) and methylation level differences in different sequence contexts under salinity stress in IR64 (E) and Pokkali (F) are shown. (G) Differential expression profiles of genes encoding DNA methyltransferases, DNA glycosylases, and genes involved in RdDM pathway and chromatin remodeling under desiccation stress in IR64 and Nagina 22, and under salinity stress in IR64 and Pokkali are shown via heatmap. Scale at the bottom represents log_2_ fold-change of expression under stress conditions.

Likewise, we analyzed DMRs under salinity stress in IR64 and Pokkali. A total of 2511 and 3580 DMRs representing 2314 and 3246 genes were detected in IR64 and Pokkali, respectively, under salinity stress (Fig. 4D). Majority of DMRs were detected in CHH context (76.65-77.46) followed by CG (17.92-18.44) and CHG (4.62-4.92) contexts in both the cultivars. In IR64, hypermethylated DMRs in all the sequence contexts; CG (63.56%), CHG (58.62%) and CHH (57.99%), were more in number under salinity stress. In contrast, hypomethylated DMRs in CG (57.73%) and CHG (63.64%) contexts were represented in higher fraction in Pokkali. However, a significant fraction of hypermethylated DMRs in CHH context (80.69%) was observed under salinity stress in Pokkali cultivar. We analysed methylation level differences under salinity stress in both the cultivars. In IR64, marginally larger fraction of DMRs was associated with hypermethylation in CG and CHG sequence contexts. While, extent of hypermethylation in CHG context was high (Fig. 4E). Interestingly, a very large fraction of DMRs showed hypermethylation in CHH context in Pokkali, even though higher number of hypomethylated DMRs was detected in CG and CHG contexts (Fig. 4F). This suggests that CHH context hypermethylation may be involved in salinity stress response in the rice cultivars.

Next, we analysed expression profiles of genes encoding DNA methyltransferases and genes involved in chromatin remodelling, RNA-dependent DNA methylation (RdDM) pathway and demethylation (Lanciano *et al*., 2017). Interestingly, lower transcript abundance of *OsDRM1a* and *OsDRM1b* methyltransferase genes (involved in methylation) and higher expression level of *OsROS1c* genes (involved in demethylation) was detected in response to desiccation stress in Nagina 22 as compared to IR64 (Fig. 4G). This suggested that passive methylation and active demethylation in CHH context might be important for desiccation stress tolerance in Nagina 22. In addition, lower expression of *CMT3* under desiccation stress may be the possible reason for hypomethylation in CHG and CG contexts in Nagina 22. *CMT3* mediated methylation in CHG context and further its role in extending methylation in CG context has been reported ((Du *et al*., 2014; Trejo *et al*., 2017). Likewise, we detected higher expression of *OsDRM1* (involved in de-novo methylation in CHH context) and higher expression of genes involved in RdDM pathway, such as *OsDcl3b* and *OsRDR2*, under salinity stress in Pokkali as compared to IR64. In addition, lower expression of *OsDML3a* and *OsDML3b* involved in demethylation was detected. This suggests that RdDM pathway associated hypermethylation in CHH context and less demethylation may be important for salinity stress tolerance in Pokkali (Fig. 4G).

Our previous analysis showed that methylation differences among the rice cultivars with contrasting responses to drought and salinity stresses were mostly found in CG context (Garg *et al*., 2015). However, most of methylation changes under stress conditions were detected in CHH context in this study. Methylation differences in CG context between cultivars and methylation changes in CHH context within a cultivar under stress have been reported in previous studies too (Liang D *et al*., 2014; Garg *et al*., 2015; Lu *et al*., 2017). This suggests that methylation dynamics in CHH context guided by de-novo methylation and demethylation may be important during stress response in a cultivar. However, methylation differences detected in CG context between cultivars may be due to diversification of DNA methylomes during selection. Significant methylation in CG context within gene body has been detected in most of the higher plants, but not in lower organisms (Zemach *et al*., 2010b; Bewick *et al*., 2016). However, lack of gene body methylation in CG context in an angiosperm (*Eutrema salsugineum*) was found to be due to loss of CMT3 (Bewick *et al*., 2016). This suggests that methylation differences detected in CG context between cultivars may be guided by CMT3 in cultivar specific-manner. Interestingly, epigenetic memory was found to be transmitted through DNA methylation in CG context in subsequent generations (Mathieu *et al*., 2007; Reinders *et al*., 2009). It is possible that epigenetic memory in subsequent exposures within a generation may be retained through DNA methylation in CHH context.

Next, we analysed gene ontology (GO) terms represented in hyper/hypomethylated genes under stress conditions. In Nagina 22, GO terms associated with abiotic stress response, including response to desiccation, were enriched in hypomethylated genes. In Pokkali, GO terms related to abiotic stress response, including response to salt stress in the hypermethylated genes (Fig. S4). These results suggest that hypomethylation under desiccation stress in Nagina 22 and hypermethylation under salinity stress in Pokkali may be important to elicit stress tolerance in these cultivars.

### Correlation between differential methylation and differential gene expression under desiccation stress

To further understand the role of DNA methylation in response to desiccation stress, we analyzed differential gene expression of DMR associated genes in IR64 and Nagina 22 cultivars under desiccation stress. In IR64, a total of 163 DMR-associated genes exhibited differential expression under desiccation stress (Fig. 5A). Interestingly, 84.1% of these genes were associated with DMRs in CHH context (Fig. 5C). In Nagina 22, a total of 178 DMR-associated differentially expressed genes were identified and most (87.77%) of these genes showed differential methylation in CHH context (Fig. 5B, D). These results suggest that methylation dynamics in CHH context is involved in desiccation stress response in both the cultivars. Next, we analyzed the DMRs found in CHH context in different gene regions. Interestingly, most (83-86.5%) of the DMRs were located in flanking regions in both the cultivars (Fig. 5B, D). Majority (69.1-71%) of the CHH context DMR-associated genes showed higher expression in both the cultivars under desiccation stress (Fig. 5B, D). However, a negative correlation of hypomethylation with higher gene expression was observed only in Nagina 22. In IR64, about 45-50% of the genes showing hypomethylation in CHH context in different gene regions exhibited higher expression under desiccation stress, suggesting no obvious correlation between differential methylation and differential gene expression (Fig. 5B). In contrast, about 77-84% of the genes that showed hypomethylation in CHH context in different gene regions exhibited enhanced transcript abundance under desiccation stress in Nagina 22 (Fig. 5D).

**Fig. 5.**
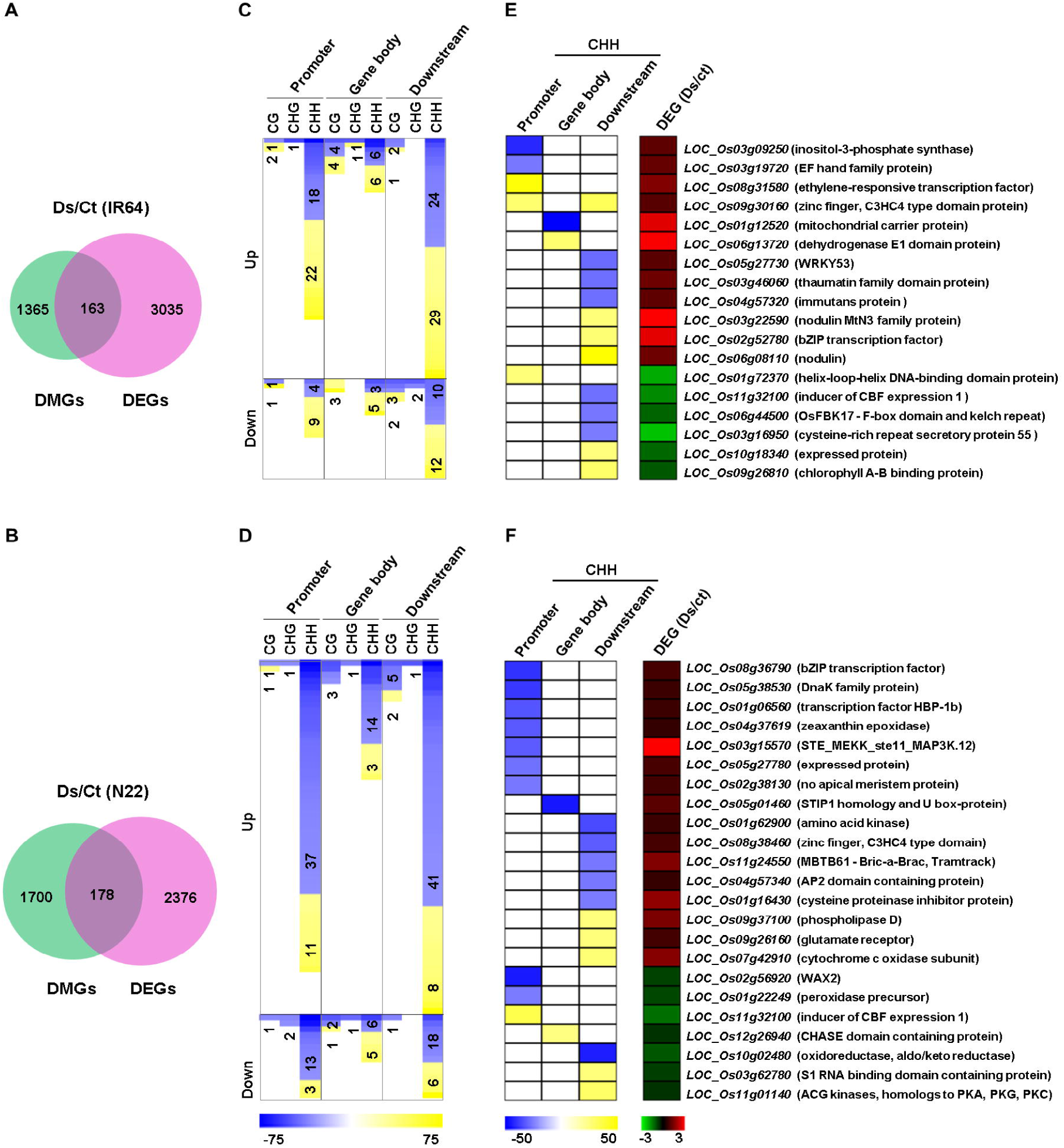
Differential methylation associated with desiccation stress response in rice. (A, B) Number of DMR associated differentially expressed genes under desiccation stress in IR64 (A) and Nagina 22 (B) is given. (C, D) Differential methylation profiles of the up/down-regulated genes in different sequence contexts and gene regions in IR64 (C) and Nagina 22 (D) are shown. (E, F) Differential methylation (CHH context) profiles of up/down-regulated genes involved in abiotic stress response in IR64 (E) and Nagina 22 (F) are shown. The right most heatmaps show differential gene expression profiles under the stress conditions in IR64 (E) and N22 (F). Scales at the bottom represent methylation level difference (blue, hypomethylation; yellow, hypermethylation) and expression log_2_ fold-change (red, upregulation; green, downregulation) under stress condition.

Next, we analyzed methylation changes under desiccation stress in sets of genes known to be involved in abiotic stress response. In IR64, a total of 18 DMR-associated genes involved in abiotic stress response showed differential expression under desiccation stress. Of these, 12 genes showed enhanced expression and 50% of them were either hyper/hypomethylated under desiccation stress, suggesting no obvious correlation (Fig. 5C). In contrast, 13 genes representing 81.25% of the CHH context DMR-associated genes exhibited hypomethylation and higher gene expression under desiccation stress in Nagina 22 (Fig. 5F). Hypomethylation was found mostly in flanking regions (92.3%) for this set of genes. Genes encoding transcription factor(s) and/or those involved in abiotic stress response, including bZIP (*LOC_Os08g36790*), zinc finger (*LOC_Os08g38460*), AP2 (*LOC_Os04g57340*), no apical meristem (*LOC_Os02g38130*), and homeobox (*LOC_Os01g06560*), were found to be correlated with hypomethylation and higher gene expression under desiccation stress in Nagina 22. Gain of function of bZIP (*OsbZIP71*), zinc finger (*OsSAP1*), a member of APETELLA 2 (*OsAP37*) and no apical meristem (NAM) transcription factors conferred tolerance to desiccation stress in rice (Mukhopadhyay *et al*., 2004; Oh *et al*., 2009; Liu *et al*., 2014; Rahman *et al*., 2016). A gene involved in photochemical quenching and dissipation of excess light, zeaxanthin epoxidase gene (*LOC_Os04g37619*), showed hypomethylation in CHH context and higher expression. Zeaxanthin epoxidase (ZEP) gene(s) are known to involved in abscisic acid (ABA) synthesis pathway and over-expression of an *AtZEP* gene elicited tolerance to drought stress in Arabidopsis (Park *et al*., 2008). Other genes involved in abiotic stress responses, such as cysteine proteinase inhibitor (*LOC_Os01g16430*), phospholipase (*LOC_Os09g37100*), and cytochrome c oxidase (*LOC_Os07g42910*) too showed hypomethylation and higher expression under desiccation stress. It has been shown that over-expression of chymotrypsin inhibitor-like 1 (*OsCPI1*) in rice resulted in better survival and higher seed yield under severe drought (Huang *et al*., 2007). The role of phospholipase in eliciting drought tolerance via modulating calcium-signalling pathway has also been demonstrated in rice (Abreu *et al*., 2018; Deng *et al*., 2019). These results suggest that hypomethylation in CHH context may govern regulation of candidate genes involved in desiccation stress response in Nagina 22.

### Correlation between differential methylation and differential gene expression under salinity stress

To understand the role of DNA methylation in response to salinity stress, we analyzed differential expression of the DMR-associated genes in different sequence contexts and gene regions in IR64 and Pokkali. A total of 39 and 80 DMR-associated differentially expressed genes under salinity stress were detected in IR64 and Pokkali, respectively (Fig. 6A, B). Most (86.8-90%) of these genes showed differential methylation in CHH context in both the cultivars (Fig. 6C, D). About (86.8-92%) of the DMR-associated genes in CHH context showed higher expression under salinity stress in both the cultivars and most of these genes harbored DMRs in their flanking regions (81.9-86.1%).

**Fig. 6.**
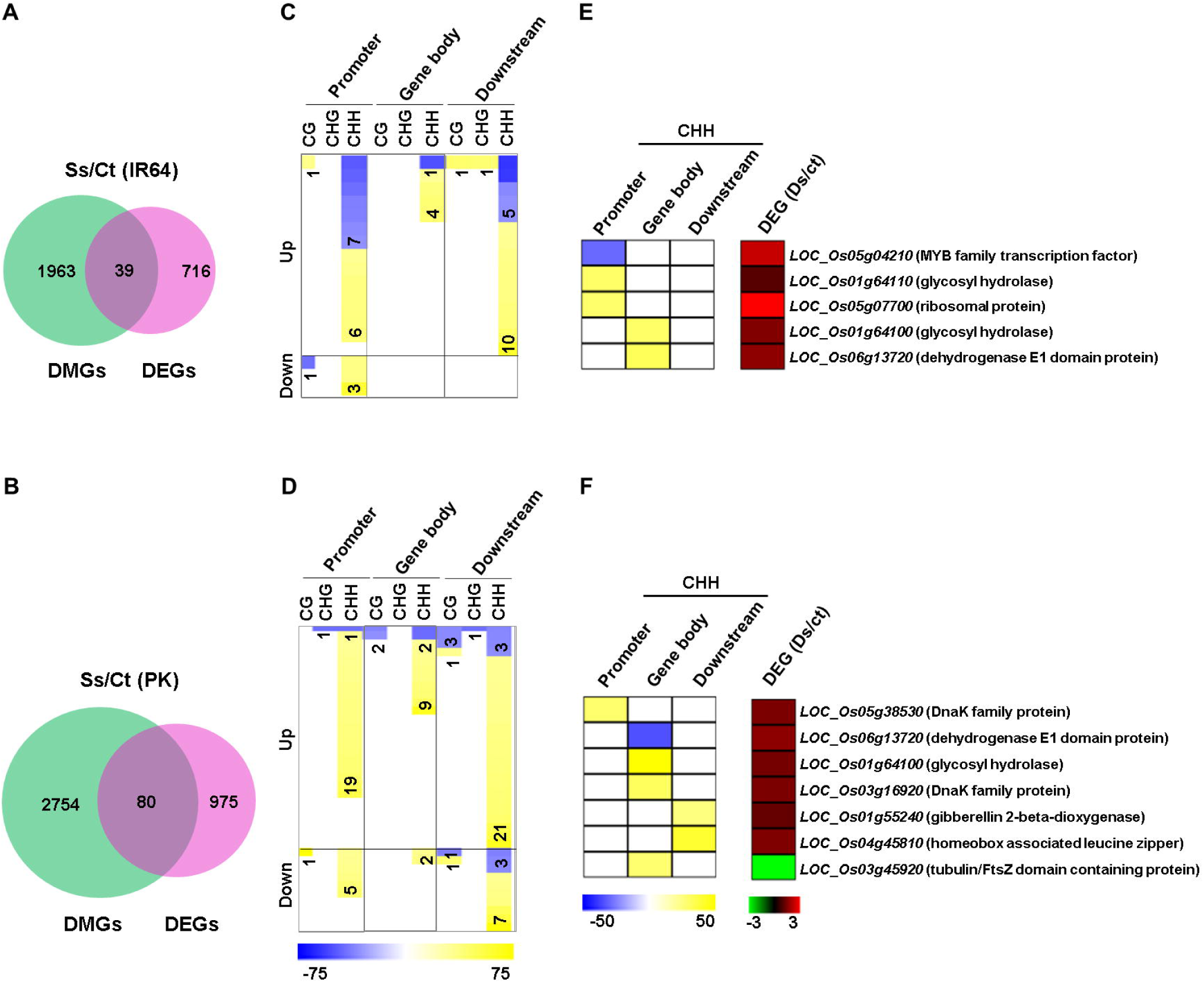
Differential methylation associated with salinity stress response in rice. (A, B) Number of DMR associated differentially expressed genes under desiccation stress in IR64 (A) and Pokkali (B) is given. (C, D) Differential methylation profiles of the up/down-regulated genes in different sequence contexts and gene regions in IR64 (C) and Pokkali (D) are shown. (E, F) Differential methylation (CHH context) profiles of the up/down-regulated genes involved in abiotic stress response in IR64 (E) and Pokkali (F) is shown. The right most heatmaps in show differential gene expression profiles under the stress conditions in IR64 (E) and Pokkali (F). Scales at the bottom represent methylation level difference (blue, hypomethylation; yellow, hypermethylation) and expression log_2_ fold-change (red, upregulation; green, downregulation) under stress condition.

In IR64, CHH context hypermethylation in gene body (80%) and downstream regions (66.6%) was correlated with higher gene expression under salinity stress (Fig. 6B). In Pokkali, correlation of CHH context hypermethylation and higher gene expression was observed with greater significance in all the gene regions, such as promoter (95%), gene body (81.8%) and downstream (87.5%) regions (Fig 6E). In addition, the number of genes showing correlation was much higher (∼5 times) in Pokkali as compared to IR64. These results suggest that hypermethylation in CHH context may be associated with salinity stress response in Pokkali. Further, we analyzed correlation of differential methylation with differential gene expression in the sets of genes involved in abiotic stress response. In IR64, four such genes exhibited CHH context hypermethylation and higher expression under salinity stress (Fig. 6E). In Pokkali, a total of five genes involved in abiotic stress response showed hypermethylation in CHH context and higher expression under salinity stress (Fig. 6F). A gene encoding transcription factor, homeobox/leucine zipper (*LOC_Os04g45810*) and other genes involved in abiotic stress response, including dehydrogenase (*LOC_Os06g13720*) and glycosyl hydrolase (*LOC_Os01g64100*) showed hypermethylation in CHH context and higher expression under salinity stress in Pokkali (Fig. 6F). Positive regulation of salinity stress response genes via leucine zipper (bZIP) has been revealed in earlier studies (Xiang *et al*., 2008; Zhao *et al*., 2014). Aldehyde dehydrogenase gene(s) play a major role in betaine biosynthesis and were found to be associated with regulation of cellular osmolytes in eliciting salinity tolerance in Arabidopsis and rice (Kishitani *et al*., 2000; Sunkar *et al*., 2003). A glycosyl hydrolase gene of rice (*OsGH5BG*) was also found to be induced in response to salinity stress (Opassiri *et al*., 2007). These results suggest that candidate genes are activated under salinity stress in the rice cultivars.

This is interesting that desiccation stress response in Nagina 22 is mediated via hypomethylation in CHH context and salinity stress response in Pokkali is mediated via hypermethylation in CHH context. Previous studies have also demonstrated that hypomethylation associated with higher gene expression under drought/salinity stress response and hypermethylation associated with higher gene expression under salinity stress response (Boyko *et al*., 2010; Wang *et al*., 2014; Al-Lawati *et al*., 2016). These results suggest that abiotic stress response(s) is dependent on cultivar specific manner.

## Conclusions

In this study, we showed an important role of DNA methylation in abiotic stress responses in desiccation tolerant (Nagina), salinity tolerant (Pokkali) and sensitive (IR64) rice cultivars. Methylation in CHH context was found to be most dynamic under desiccation and/or salinity stress conditions in all the rice cultivars. Interestingly, hypomethylation in CHH context was correlated with higher gene expression under desiccation stress in Nagina 22. In contrast, hypermethylation in CHH context was correlated with higher gene expression under salinity stress in Pokkali. These results revealed that abiotic stress response is cultivar-specific in rice. Altogether, we provided new insights into role of DNA methylation in response to abiotic stress in rice.

## Supplementary data

Supplementary data are available at *JXB* online.

**Table S1.** Summary of bisulphite sequencing data, mapping and methylated cytosines.

**Fig. S1.** Average methylation level in different sequence contexts under control and stress conditions.

**Fig. S2.** Methylation level in forward and reverse strands.

**Fig. S3.** Methylation patterns in CHH context in TEs.

**Fig. S4.** Gene ontology (GO) analysis of differentially methylated genes under abiotic stress conditions.

## Acknowledgements

This work was financially supported by the Jawaharlal Nehru University (JNU), New Delhi under the University with Potential for Excellence (UPoE II) scheme of the University Grants Commission, New Delhi. We are thankful to Mr. VVS Narayana Chevala for help with handling methylation data at initial steps.

